# Endogenous retroviruses drive species-specific germline transcriptomes in mammals

**DOI:** 10.1101/2020.03.11.987230

**Authors:** Akihiko Sakashita, So Maezawa, Kris G. Alavattam, Masashi Yukawa, Artem Barski, Mihaela Pavlicev, Satoshi H. Namekawa

## Abstract

Gene regulation in the germline ensures the production of high-quality gametes, long-term maintenance of the species, and speciation. Germline transcriptomes undergo dynamic changes after the mitosis-to-meiosis transition in males and have been subject to evolutionary divergence among mammals. However, the mechanism that underlies germline regulatory divergence remains undetermined. Here, we show that endogenous retroviruses influence species-specific germline transcriptomes in mammals. We show that the expression of endogenous retroviruses, particularly the evolutionarily young K family (ERVK), is associated with gene activation after the mitosis-to-meiosis transition in male mice. We demonstrate that accessible chromatin and H3K27ac, a marker of active enhancers, are tightly associated with ERVK loci as well as with the activation of neighboring evolutionarily young germline genes. Thus, ERVKs serve as evolutionarily novel enhancers in mouse spermatogenesis. These ERVK loci bear binding motifs for critical regulators of spermatogenesis such as A-MYB. The genome-wide transposition of ERVKs might have rewired germline gene expression in a species-specific manner. Notably, these features are present in human spermatogenesis, but independently evolved ERVs are associated with expression of germline genes, demonstrating the prevalence of ERV-driven mechanisms in mammals. Together, we propose a model whereby species-specific transcriptomes are fine-tuned by endogenous retroviruses in the mammalian germline.

## Introduction

The testis has the most diverse, complex, and rapidly-evolving transcriptome as well as the largest number of specifically expressed transcription factors (TFs) of all organs in mammals^1-4^. This is due, in part, to the specific burst of expression of thousands of germline genes after the mitosis-to-meiosis transition^5,6^. Meiosis is an essential process in the preparation of haploid gametes, and the mitosis-to-meiosis transition is a key switch for the genome. Notably, a wide variety of species-specific transcripts have been identified in late spermatogenesis^3,7^, giving rise to morphologically and functionally diverse gametes in mammals. However, the mechanisms that enable the rapid evolution of species-specific germline transcriptomes remain to be determined. In this study, we identify a mechanism that underlies germline regulatory divergence. We report that many rapidly-evolved cis-regulatory elements, specifically active enhancers, are derived from a group of endogenous retroviruses (ERVs). ERVs are the remnants of retroviruses that integrated into the germline genome. A large group of ERVs, long terminal repeat (LTR)-type transposable elements (TEs), are recognizable due to a specific portion of the viral sequence and constitute approximately 10% of mammalian genomes^8^.

TEs are mobile genetic elements that account for approximately 40-50% of a given mammalian genome^9^. TEs are comprised of sub-categories of elements, such as long and short interspersed nuclear elements (respectively, LINEs and SINEs), and were considered genetic threats due to their copy-and-paste mechanisms of transposition, which are kept in check through strict silencing mechanisms. In contrast, the geneticist Barbara McClintock, who originally discovered TEs, proposed in 1950 that TEs function as gene regulatory elements^10^. Studies in the last decade, long after McClintock’s proposal, have indeed established that TEs can impact host genomes by introducing gene regulatory elements, such as promoters and enhancers^11-15^. While many TEs have lost the ability to transpose themselves, TEs have nonetheless adapted to regulate gene expression in host genomes.

The mobility of TEs in the germline is tightly controlled due to their deleterious actions on the heritable genome. The germline draws on several TE-suppressing mechanisms, including DNA methylation, H3K9 methylation, and PIWI-interacting RNA (piRNA)^16-18^. Yet despite such silencing mechanisms, recent studies have revealed regulatory functions for TEs in male meiosis. These encompass post-transcriptional functions to regulate mRNA and long noncoding RNAs (lncRNAs) via the piRNA pathway^19^, and promoter functions for LTRs that drive the expression of lncRNAs^20^. However, the function of TEs as cis-regulatory elements remains undetermined at the mitosis-to-meiosis transition, when dynamic reorganization of 3D chromatin and the epigenome takes place^6,21-24^.

In this study, we use an unbiased, genome-wide approach to identify a group of ERVs, the evolutionarily young ERVKs, that are expressed and denoted by accessible chromatin after the mitosis-to-meiosis transition. We show that ERVKs function as species-specific enhancers in the germline, providing binding sites for A-MYB (also known as MYBL1), a key transcription factor for germline genes^25,26^. These enhancers drive expression of evolutionarily novel germline genes after the mitosis-to-meiosis transition, thereby defining the species-specificity of germline transcriptomes in mammals. Notably, we demonstrate the prevalence of ERV-driven germline genes in humans. We propose a model whereby ERVs fine-tune species-specific transcriptomes in the mammalian germline.

## Results

### A large group of endogenous retroviruses are differentially expressed at the mitosis-to-meiosis transition in spermatogenesis

To determine the expression dynamics of repetitive elements interspersed throughout the genome in spermatogenesis, we analyzed previously published RNA-seq datasets for representative stages of spermatogenesis^5,6,27^ using RepBase, a database of consensus sequences for repetitive DNA elements^28^. We included in the analysis the transcriptomes of THY1^+^ undifferentiated spermatogonia from postnatal day 7 (P7) testes, which contain spermatogonial stem cells and progenitor cells; KIT^+^ differentiating spermatogonia from P7 testes; purified pachytene spermatocytes (PS) in the midst of meiosis; postmeiotic round spermatids (RS) from adult testes; and mature sperm (Fig. 1a). Among the 1,195 consensus repetitive DNA elements of *Mus musculus* in RepBase, we detected 307 differentially expressed types of repetitive elements between at least two stages of spermatogenesis (Fig. 1b, Supplementary Table 1; differentially expressed repetitive elements were defined as those with a >2-fold change using the software edgeR^29^; false discovery rate (FDR) < 0.05). Among the 307 repetitive elements, we identified two major classes of expressed repetitive elements by k-means clustering: One comprises a “mitotic type” of elements that is active in mitotic spermatogonia but repressed after meiosis (25.7%, 79/307); the other comprises a “meiotic type” that is expressed specifically after the mitosis-to-meiosis transition (74.2%, 228/307; Fig. 1b). These results reveal a large group of repetitive elements that are highly expressed after the mitosis-to-meiosis transition. Interestingly, in both classes, more than half of the expressed repetitive elements belong to ERVs, the type of TE characterized by transcription factor binding site-rich long terminal repeats (LTR) (Fig. 1c).

**Figure 1.**
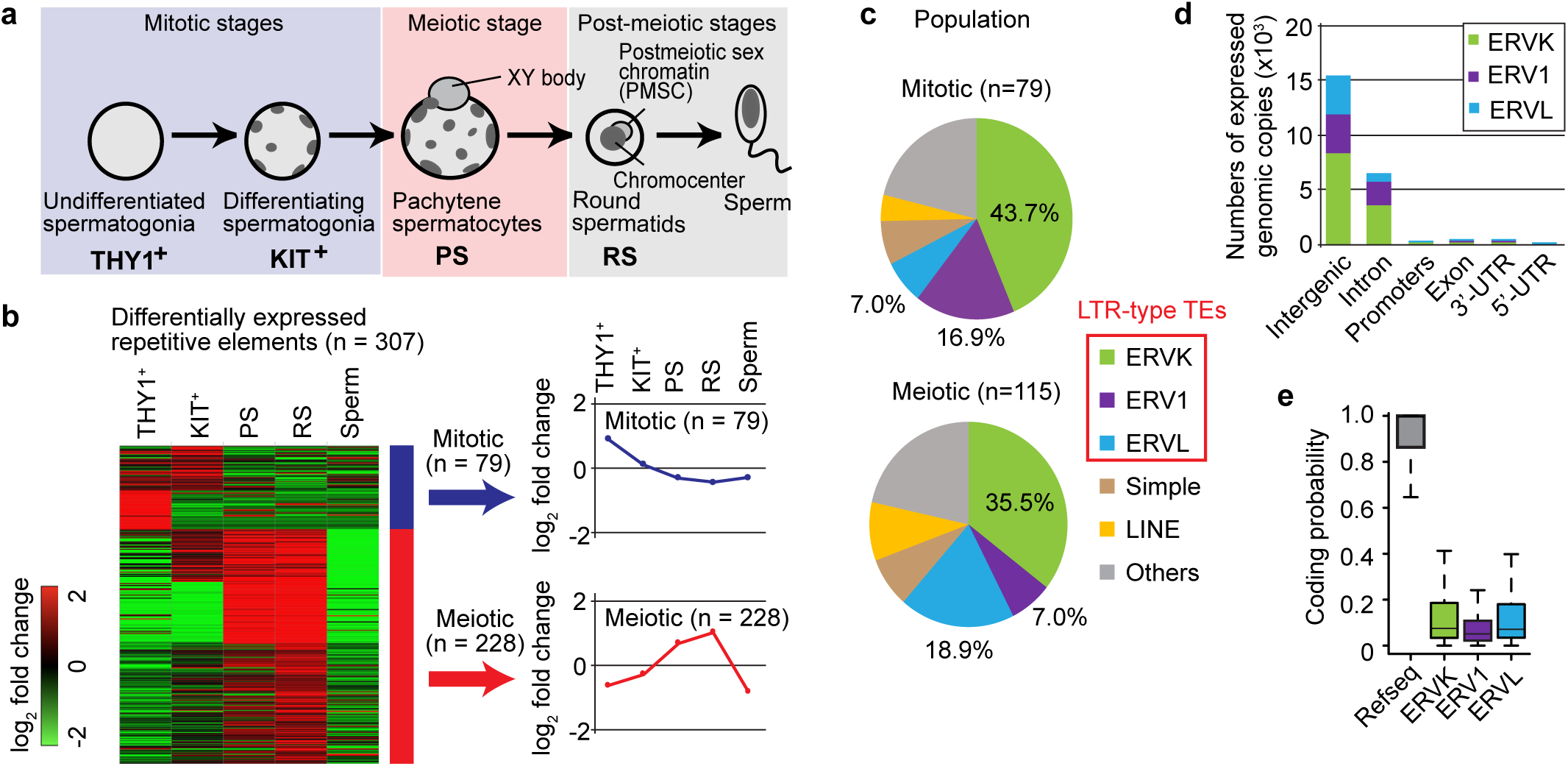
Dynamic expression of repetitive elements during mouse spermatogenesis. (**a**) Schematic of mouse spermatogenesis and the five representative stages analyzed in this study: THY1^+^, undifferentiated spermatogonia; KIT^+^, differentiating spermatogonia; PS, pachytene spermatocyte; RS, round spermatids; Sperm: epididymal spermatozoa. (**b**) Differentially expressed repetitive elements were defined as those with a >2-fold change using the software edgeR; false discovery rate (FDR) < 0.05. A total of 307 differentially expressed repetitive elements were classified into two clusters. Right panels show mean values of log_2_ fold changes in each cluster. (**c**) Pie charts indicating the population of the major classes of repetitive elements in each cluster. (**d**) Enrichment of ERVs expressed in each genomic region. (**e**) Box and whisker plots showing the protein coding probabilities (calculated with CPAT; see Methods) of ERV1, ERVK, and ERVL loci in comparison to RefSeq genes. Central bars represent medians, the boxes encompass 50% of the data points, and the error bars indicate 90% of the data points.

Next, to determine the genomic locations of elements for each expressed ERV family, we aligned the RNA-seq data of representative stages of spermatogenesis to the unique annotated ERV sequences in the mouse genome (mm10, Genome Reference Consortium Mouse Build 38 (GRCm38)). A vast majority of ERVs expressed in PS mapped to intergenic and intronic regions (Fig. 1d). Interspersed ERVs code for a series of proteins (*gag, pro, pol*, and *env*) flanked by LTRs that are required for copy number expansion in a host genome^30^; to determine the extent of protein coding by ERVs after the mitosis-to-meiosis transition, we predicted coding probabilities using the bioinformatic tool Coding Potential Assessment Tool (CPAT; see Methods)^31^. We observed that protein-coding ERVs were significantly lower than predicted and, indeed, lower than that of reference sequence (RefSeq) genes, including coding and non-coding genes (Fig. 1e). These results suggest that expressed ERVs do not function as coding proteins and may have lost their transposon activities. Therefore, we infer that ERVs expressed in mouse spermatogenesis have functions other than coding proteins.

### Accessible chromatin is uniquely established at a subset of ERVs in late spermatogenesis

Since our expression analyses suggested that a large number of ERVs are differentially expressed in spermatogenesis and unlikely to have transposon activity, we sought to determine the accessibility of chromatin for these ERVs in spermatogenesis, since accessible chromatin is a notable feature of gene regulatory elements. To this end, we analyzed previously published ATAC (assay for transposase-accessible chromatin using sequencing)-seq datasets^21,32^ that contain the sites of accessible chromatin during spermatogenesis. Genome-wide, we extracted 733,999 unique ERV loci, including all subfamilies of ERV1, ERVK, and ERVL, per UCSC RepeatMasker annotation, and aligned the ATAC-seq data to these ERV loci. While we found that most ERV loci evince closed chromatin (Supplementary Fig. 1), we detected a subset of genomic ERV loci that evince changes in chromatin accessibility during spermatogenesis (Fig. 2a). Through k-means clustering, we identified two major classes of accessible ERV loci: One comprises a mitotic type, composed of loci that are accessible in mitotic spermatogonia but closed after meiosis (consisting of 69.1% of accessible ERV1s, 5.8% of accessible ERVKs, and 34.4% of accessible ERVLs); the other comprises a meiotic type, composed of chromatin loci that are open in PS and RS, having become accessible after the mitosis-to-meiosis transition (consisting of 30.9% of accessible ERV1s, 94.2% of accessible ERVKs, and 65.6% of accessible ERVLs; Fig. 2a). Of note, most differentially accessible ERVK loci were classified into the meiotic type, and approximately 87% of accessible ERVKs in meiosis were members of one of three subgroups: RLTR10 (65.7%, 394/600), RMER17 (11.5%, 69/600), or RLTR51 (9.5%, 57/600; Fig. 2b). We also detected RLTR23, RLTR41, and RMER5 as the major subgroups of meiotic ERV1 loci; ORR1, MTE, and MTE2 comprised the major subgroups of meiotic ERVL loci. Together, we identified groups of ERVs that become accessible during meiosis, with a notable enrichment of ERVK loci, especially RLTR10, RMER17, and RLTR51.

**Figure 2.**
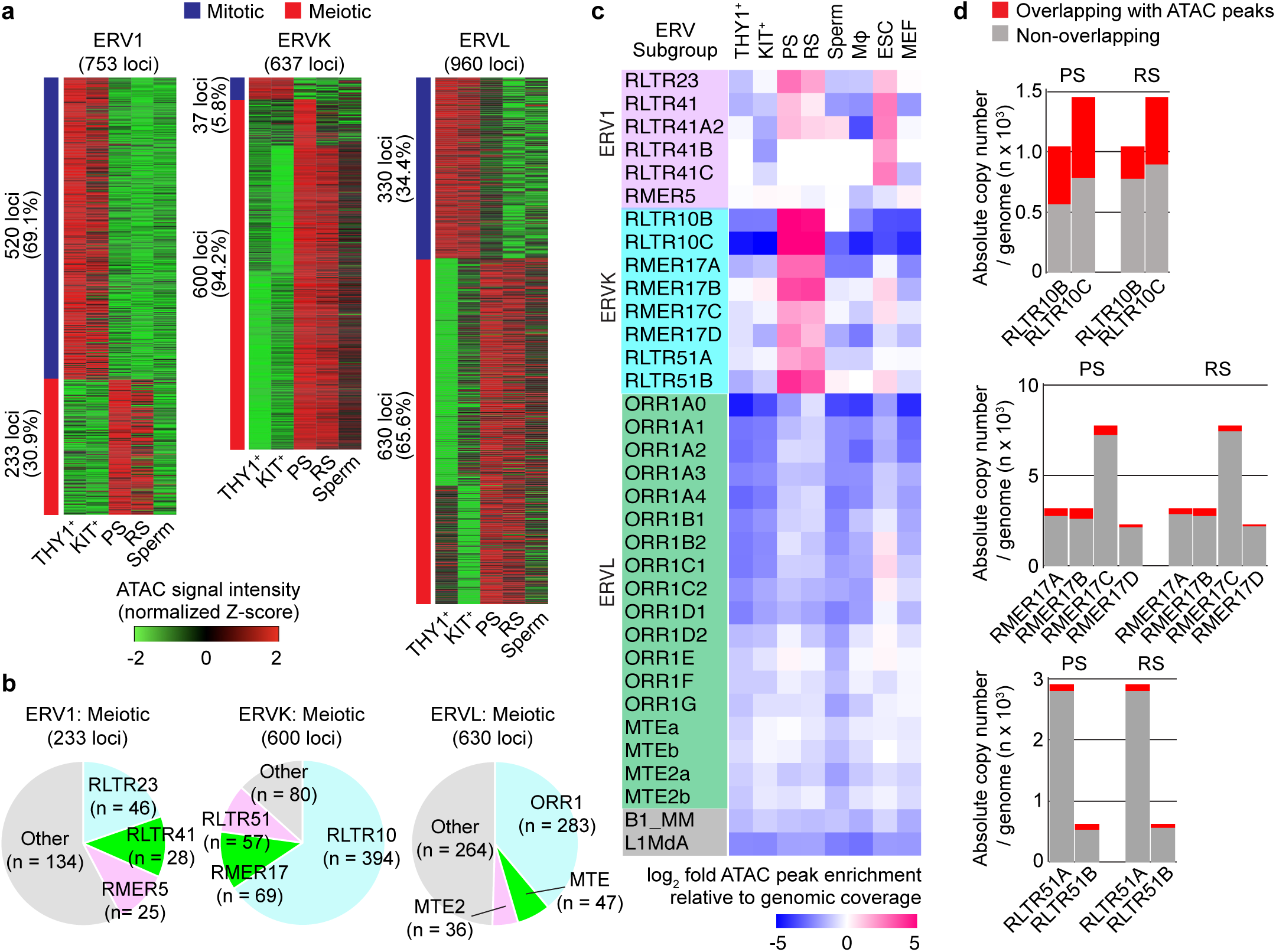
Evolutionary young ERVKs uniquely form accessible chromatin in late spermatogenesis. (**a**) Heat maps depicting changes in the chromatin accessibility of ERVs interspersed throughout the genome, having been identified with the program edgeR (>2-fold change, Bonferroni: *P* < 0.05, binomial test; see Methods). (**b**) Pie charts indicating the absolute abundance of respective ERV subgroups comprising the meiotic cluster. (**c**) Heat maps depicting log_2_ fold enrichment of specific classes of ERVKs in ATAC-seq-enriched regions relative to genomic prevalence; ATAC-seq-enriched regions were identified with the program MACS (see Methods). (**d**) Bar charts indicating the absolute copy numbers of ERVK subgroups non-overlapping (gray) and overlapping (red) ATAC-seq- enriched regions.

To confirm that the enrichment of accessible chromatin at ERV loci is specific to late spermatogenesis, we examined the enrichment of ATAC-peaks for each subgroup of ERV subfamilies in other tissues (Fig. 2c). For this analysis, we examined ATAC-seq data from lung macrophages (MΦ), embryonic stem cells (ESCs), and embryonic fibroblasts (MEF) as controls. Of note, we found that accessible chromatin was specifically enriched for each subgroup of evolutionarily young ERVKs: RLTR10 (RLTR10B and RLTR10C), RMER17 (RMER17A, RMER17B, RMER17C, and RMER17D), and RLTR51 (RLTR51A and RLTR51B) (Fig. 2c). Furthermore, among ERV1s, accessible chromatin was also enriched at RLTR23, RLTR41, and RLTR41A2 loci in PS and RS. Curiously, we also observed an enrichment of accessible chromatin at several subgroups of ERV1 in ESCs. By contrast, enrichment of accessible chromatin was not observed at ERVLs, SINEs (B1_MM), and LINEs (L1MdA; Fig. 2c). Since chromatin accessibility was the highest at ERVK loci, we hereafter focus on the analyses of these elements.

Notably, among genomic ERVK loci, nearly half of RLTR10B and RLTR10C overlapped accessible chromatin in PS and RS (Fig. 2d). Although we also observed the enrichment of accessible chromatin on subgroups of RMER17 and RLTR51 in comparison to average genomic enrichment (Fig. 2c), only a relatively minor population of these loci overlap with accessible chromatin (Fig. 2d). Therefore, RLTR10B and RLTR10C are of particular interest in the context of late spermatogenesis. We further identified three common features of ERVs overlapping accessible chromatin during meiosis. First, accessible ERVK loci largely overlap between PS and RS (Supplementary Fig. 2a). Second, these loci are largely located in intergenic regions, while only a minor portion is located in intronic regions (Supplementary Fig. 2b). Third, accessible ERVKs, particularly RLTR10 and RMER17, were detected on the sex chromosomes with much higher frequency than on autosomes in PS (Supplementary Fig. 2c). In PS, the sex chromosomes undergo a tightly coordinated process of transcriptional inactivation known as meiotic sex chromosomes inactivation (MSCI); it is in this context that, counterintuitively, the accessibility of chromatin increases^21^. Our data suggest that subgroups of RLTR10 and RMER17 become accessible despite MSCI. Taken together, these data demonstrate that specific types of ERVs, especially evolutionarily young ERVKs, form accessible chromatin in late spermatogenesis.

### ERVKs that are accessible in late spermatogenesis have enhancer-like features

Since ERVKs accessible in late spermatogenesis are largely located in intergenic regions, we hypothesized that they function as cis-regulatory elements, such as enhancers, that drive the expression of spermatogenesis-specific genes. To test this hypothesis, we examined the distribution of three histone modification marks around accessible ERVK loci: active histone modifications representing (1) active enhancers, H3K27ac, and (2) promoters, H3K4me3, as well as a repressive histone modification representing (3) facultative heterochromatin, H3K27me3. We analyzed the average tag density of chromatin immunoprecipitation sequencing (ChIP-seq) data^33,34^ (Maezawa et al., co-submitted) for these modifications in regions surrounding ± 1 kb of accessible ERVKs in representative stages of spermatogenesis. Notably, H3K27ac was significantly enriched within accessible ERVK loci in PS and RS (Fig. 3a), suggesting that accessible ERVKs evince features of active enhancers in late spermatogenesis. On the other hand, throughout spermatogenesis, profiles for H3K4me3 and H3K27me3 did not manifest dynamic changes at accessible ERVKs (Fig. 3a), providing further evidence that ERVKs have enhancer-like features. We sought to corroborate these data through analyses of representative loci: The specific establishment of H3K27ac and open chromatin at an autosomal RTLR10 locus in PS was correlated with PS-specific upregulation of neighboring transcripts (Fig. 3b). On the X chromosome, which undergoes MSCI in PS, the establishment of H3K27ac and open chromatin at RTLR10 loci in PS correlates with activation of transcripts that escape postmeiotic silencing in RS (Fig. 3c). This establishment of H3K27ac on the silent X chromosome in meiosis is regulated by RNF8, a DNA damage response factor^34^. Therefore, on the X chromosome, ERVK loci are regulated downstream of RNF8. Importantly, both autosomal and X chromosomal RTLR10 loci do not overlap with transcription start sites (Fig. 3b and 3c), suggesting that RTLR10 loci have functions akin to enhancers rather than promoters. Strikingly, this is in contrast to the proposed functions of ERVs in spermatogenesis, in which ERVs act as promoters to drive the expression of lncRNAs in late spermatogenesis^20^. Taken together, we found that accessible ERVKs are enriched with H3K27ac and are likely to act as enhancers. A previous study demonstrated that, in placenta, one ERV subgroup, RLTR13D5, functions as a species-specific enhancer, suggesting that ERVs may have enhancer functions in other tissues, such as testes, as well^35^. Our results corroborate this notion: Specific subgroups of ERVKs gain features of active enhancers in late spermatogenesis.

**Figure 3.**
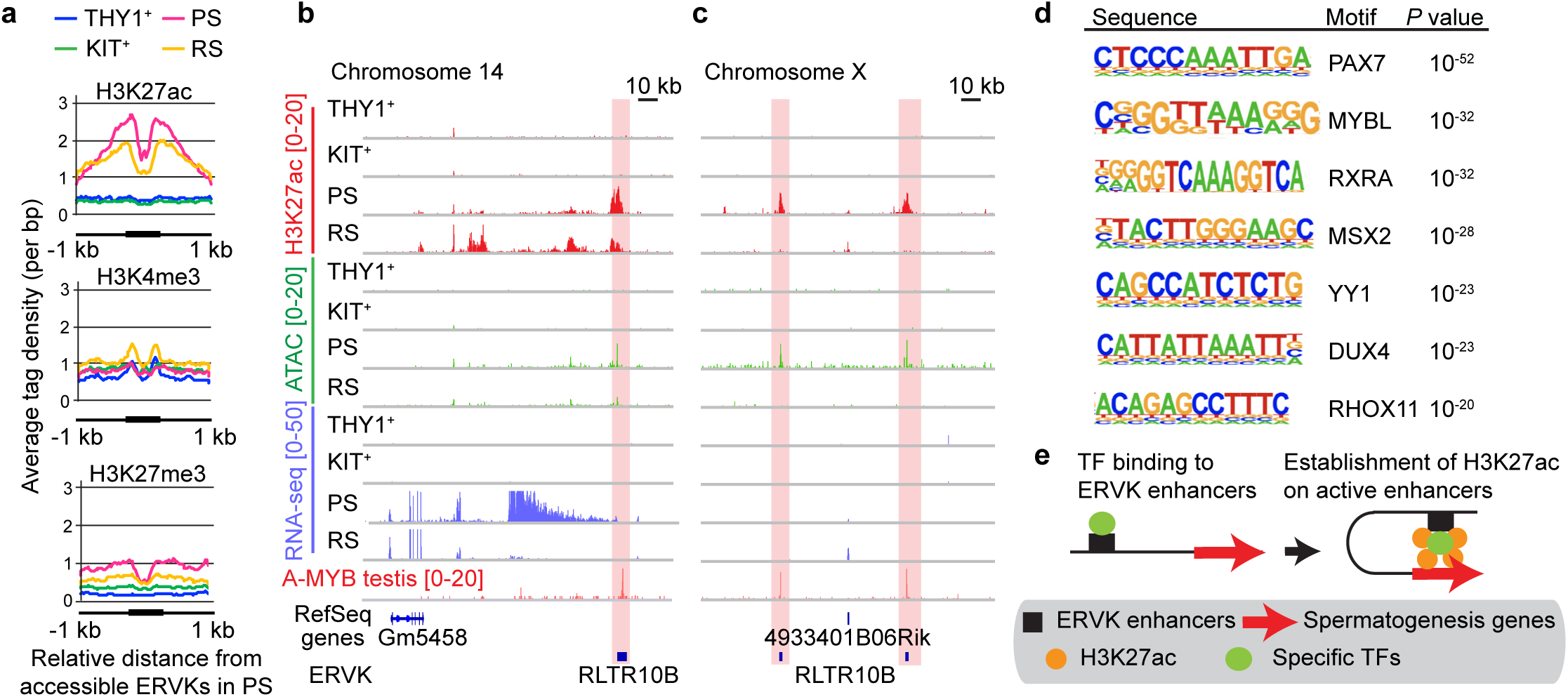
Young ERVKs at accessible chromatin harbor active enhancer marks and transcription factor binding sites. (**a**) Average tag density plots of H3K27ac, H3K27me3, and H3K4me3 ChIP-seq signals around accessible ERVKs (± 1 kb) in pachytene spermatocytes (PS). The active enhancer mark H3K27ac is significantly enriched at accessible ERVK loci. (**b, c**) Track views showing H3K27ac ChIP-seq, ATAC-seq, RNA-seq, and A-MYB ChIP-seq signals on chromosome 14 (autosome example, **b**) and chromosome X (sex chromosome example, **c**) in spermatogenic cells. Red rectangles indicate accessible ERVK loci as denoted by H3K27ac ChIP-seq, ATAC-seq, and A-MYB ChIP-seq signals. (**d**) Motif analyses of ERVKs at accessible chromatin for putative transcription factor (TF)-binding sites as called by HOMER (see Methods). (**e**) Model for young ERVK enhancers as activators of germline genes.

ERVs are known to carry binding sites for TFs and, therefore, bear the potential to rewire transcriptomes via transposition^11-15^. To determine the TF binding sites present in accessible ERVKs, we performed motif analyses using the program HOMER^36^. We identified binding sites of TFs that are implicated in spermatogenesis and meiosis (PAX7, MYBL, RXRA, MSX2, YY1, DUX4, and RHOX11: Fig. 3d)^25,26,37-41^. Importantly, recent studies demonstrated that A-MYB (also known as MYBL1), a male germline-specific transcription factor, drives spermatogenesis-related gene expression from meiotic prophase onward^25,26^. In support of motif analysis, A-MYB ChIP-seq peaks in whole testes^26^ overlapped with enhancer-like ERVK loci (RLTR10B loci) at intergenic regions both on autosomes and on the X chromosome (Fig. 3b, c). Consistent with this, a recent study demonstrated that A-MYB binds to RLTR10B^42^. Taken together, these analyses indicate that accessible ERVKs in late spermatogenesis serve as active enhancers by providing specific TF binding sites to drive the expression of spermatogenesis-specific transcripts (Fig. 3e).

### Accessible ERVKs act as enhancers to drive expression of adjacent genes in late spermatogenesis in an A-MYB-dependent manner

To further define the functions of accessible ERVKs as enhancers, we tested the hypothesis that genes adjacent to enhancer-like ERVKs evince preferential expression relative to non-adjacent genes after the mitosis-to-meiosis transition. To this end, we identified 1,874 genes that are adjacent to accessible ERVKs in PS (i.e., the genes nearest to accessible ERVKs in PS, see Methods). Among 1,874 genes, 543 genes (27.7%: Supplementary Table 2) overlapped with genes specifically activated in the mitosis-to-meiosis transition (5,873 preferentially expressed genes in meiosis were defined at the transition from KIT^+^ spermatogonia to PS based on two parameters: (1) a fold change (FC) in gene expression of ≥ 2 and (2) statistical significance determined using a Wald test with a Benjamini-Hochberg FDR of < 0.01: Fig. 4a). This association is statistically significant compared to the ratio of preferentially expressed genes in PS to all genes in the genome (5,873 preferentially expressed genes / all 22,661 RefSeq genes in the genome; *P* = 9.77 x 10^−4^, hypergeometric test). Thus, we denote these 543 genes as those near to accessible enhancer-like ERVKs (henceforth, we call these genes “ERVK-adjacent genes”). Importantly, the ERVK-adjacent genes were highly expressed in PS in comparison to other genes in the genome (Fig. 4b). We performed gene ontology (GO) analysis on this gene set and revealed that ERVK-adjacent genes comprise genes associated with chromosome segregation, chromatin silencing, and spermatogenesis (Fig. 4c). More than 30% of ERVK-adjacent genes were unannotated genes such as gene names starting with “*Gm*” and “*LOC*” or ending with “*Rik*”, such as *Gm10415, LOC666331* and *1110019D14Rik*. However, genes known to be involved in spermatogenesis were also detected as ERVK-adjacent genes including *Spata24, Nme8*, and *Zscan2*^43-45^. These results further suggest that accessible ERVKs function as enhancers to activate genes involved in spermatogenesis after the mitosis-to-meiosis transition.

**Figure 4.**
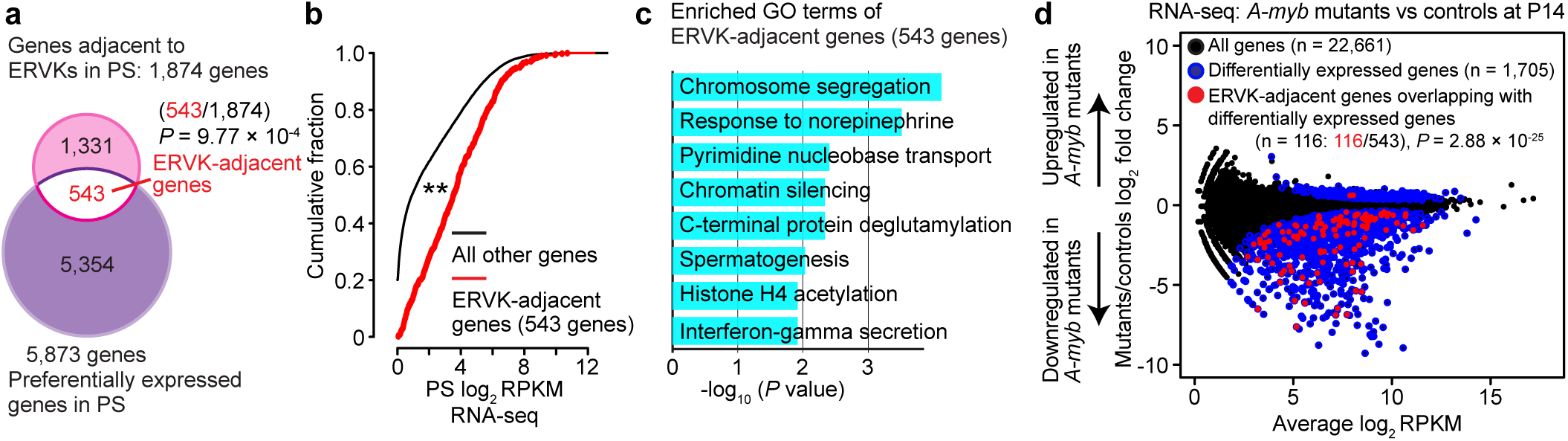
ERVKs adjacent genes are highly expressed in late spermatogenesis in an A-MYB-dependent manner. (**a**) Venn diagrams showing the overlap between genes adjacent to ERVKs in accessible chromatin (pink) and genes preferentially expressed in pachytene spermatocytes (PS; purple). *P* value is based on a hypergeometric probability test. The number of genes for *P* value are 543 genes preferentially expressed genes in PS/1,874 genes that are adjacent to accessible ERVKs in PS (*P* = 9.77× 10^−4^) compared to 5,873 preferentially expressed genes in PS/all 22,661 RefSeq genes in the genome. (**b**) A cumulative distribution plot comparing the expression of genes adjacent to ERVKs in accessible chromatin (pink) and the expression of all RefSeq genes (black) in PS. *P* value is based on a Kolmogorov-Smirnov test (**; *P* < 0.01). (**c**) Bar chart showing enriched GO terms for genes adjacent to ERVKs in accessible chromatin. (**d**) RNA-seq analysis of *A-myb* mutant versus heterozygous control testes at 14 days post-partum (dpp). The 1,705 genes evincing significant changes in expression (>2-fold change, *P adj* < 0.01: a binomial test) in *A-myb* mutants are represented by blue circles. *P* value is based on a hypergeometric probability test. The 116 dysregulated genes (represented by red circles)/543 ERVK-adjacent genes (*P* = 2.88 × 10^−25^) compared to 1,705 dysregulated genes/all 22,661 RefSeq genes in the genome.

Based on the A-MYB ChIP-seq analysis (Fig. 3b, c) and the motif analyses of enhancer-like ERVKs, we sought to test the following hypothesis: The binding of A-MYB to enhancer-like ERVKs enables activation of adjacent genes in late spermatogenesis. To that end, we analyzed previously published RNA-seq data from the testes of *A-myb* mutants (*Mybl1*^*repro9*^) and heterozygous littermate controls at P14^26^ (Fig. 4d). Consistent with the reported role of A-MYB in the activation of late spermatogenesis genes, 1,705 genes were differentially expressed and most of them were downregulated upon the loss of A-MYB (Fig. 4d). Importantly, we observed a significant overlap of ERVK-adjacent genes and genes differentially expressed in *A-myb* mutants (116 genes out of 543 adjacent genes; compared to 1,705 differentially expressed genes / all 22,661 RefSeq genes in the genome; *P* = 2.88 × 10^−25^; hypergeometric test), and many of them were found in the downregulated genes of *A-myb* mutants. Together, these results suggest that at least a portion of enhancer-like ERVKs contribute to the activation of adjacent genes through the action of A-MYB in late spermatogenesis.

### Young ERVKs regulate species-specific gene expression as enhancers during late spermatogenesis

Meiotic spermatocytes and postmeiotic spermatids manifest high levels of transcriptomic diversity across species of mammals^3,7^. Therefore, we reasoned that evolutionarily young enhancer-like ERVKs may drive newly evolved genes, thereby conferring species-specific diversity among transcriptomes during late spermatogenesis in mammals. To test this possibility, we sought to determine the degree of sequence diversity of ERVK-adjacent genes in mammals. Notably, a subset of ERVK-adjacent genes found in mice do not have unambiguous homologs in other mammals that we examined—even rat, a close clade in rodents (161/543, 29.7%; Fig. 5a). Furthermore, many ERVK-associated genes with homologs among mammals are poorly conserved, which raises the possibility of divergent functions in mouse (Fig. 5a). These results suggest that genes close to enhancer-like ERVKs are evolutionarily new in mice and/or rapidly evolved among mammals. Thus, enhancer-like ERVKs in mice are likely to regulate mouse-specific or evolutionarily diverged genes.

**Figure 5.**
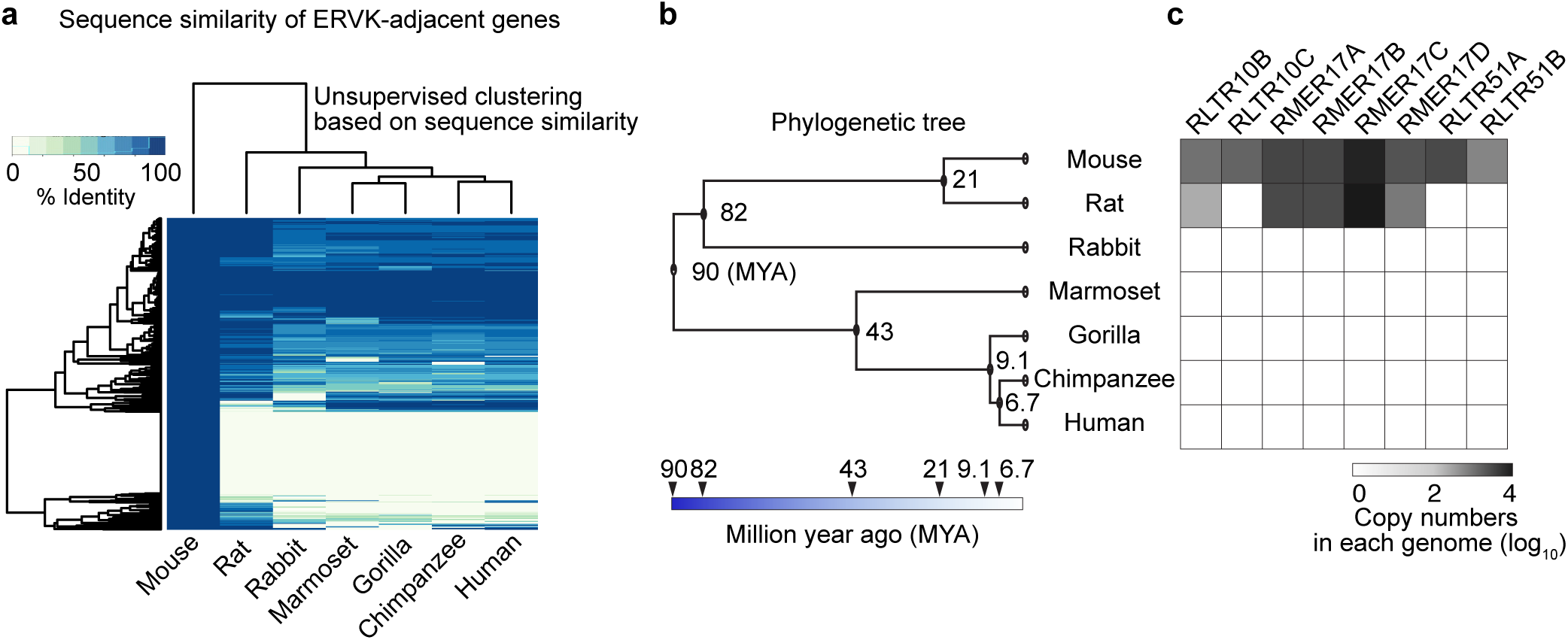
Genes adjacent to enhancer-like ERVKs are less conserved across species. Heat map showing percentage sequence identity for each of the 543 mouse genes defined as adjacent to ERVKs across 6 other species. ‘Adjacent’ genes exhibit low levels of conservation. (**b**) A phylogenetic tree indicates the evolutionary distances between rodents, rabbit and primates (adapted from TIMETREE; http://timetreebeta.igem.temple.edu/). The color scheme and respective arrowheads indicate estimated times (million years ago (MYA)) of divergence. (**c**) Heat map depicting the abundance of selected ERVK copies in respective genomes.

To determine the species-specific features of enhancer-like ERVKs, we sought to determine the evolutionary traits of young ERVKs in mammals (Fig. 5b, c). Specific members of enhancer-like ERVKs in mice are only present in rodents, and some ERVKs (RLTR10C, RLTR51A, and RLTR51B) in mice do not have counterparts even in rats (Fig. 5c). Enhancer-like ERVKs with counterparts in rats displayed varied copy numbers (Fig. 5c), and the sequences of such ERVKs are highly diverged between the two species (Supplementary Fig. 3). These results indicate that the enhancer functions of ERVKs studied here are likely specific to the regulation of spermatogenesis genes in mouse.

### A subset of ERVK and ERV1 are associated with meiotic gene expression in humans

The identification of evolutionarily young, enhancer-like ERVKs in mice prompted us to investigate the species-specific functions of ERVKs in other species of mammals. To address this, we analyzed human spermatogenesis to address the hypothesis that human-specific ERVs have enhancer-like features in human spermatogenesis. To this end, we analyzed H3K27ac ChIP-seq data from human testes. We found that MER57E3, a subgroup of ERV1, and LTR5B, a subgroup of ERVK, are enriched with H3K27ac and occupy a location adjacent to transcripts in human PS (Fig. 6a). To evaluate the genome-wide features of human ERVs, we examined the enrichment of H3K27ac on each subgroup of ERV1, ERVK, and ERVL in human testes. Consistent with our findings in mice, we detected a subset of human ERVKs that had distinctly diverged from rodent ERVKs, and this subset was enriched with H3K27ac (>2-fold enrichment compared to the expected genomic prevalence; Fig. 6b). Notably, we found a subset of human ERV1s that is highly enriched with H3K27ac (>2-fold enrichment); among them, MER57E3 showed the highest enrichment of H3K27ac (Fig. 6b). Motif analysis of human ERVs revealed that human ERV1s and ERVKs contain binding sites for A-MYB (Fig. 6c). These results suggest that, in addition to ERVKs, ERV1s act as enhancers via A-MYB-dependent mechanisms in humans. In support of this notion, we confirmed that A-MYB is highly expressed in both mouse and human spermatocytes by immunostaining testicular sections (Supplementary Fig. 4).

**Figure 6.**
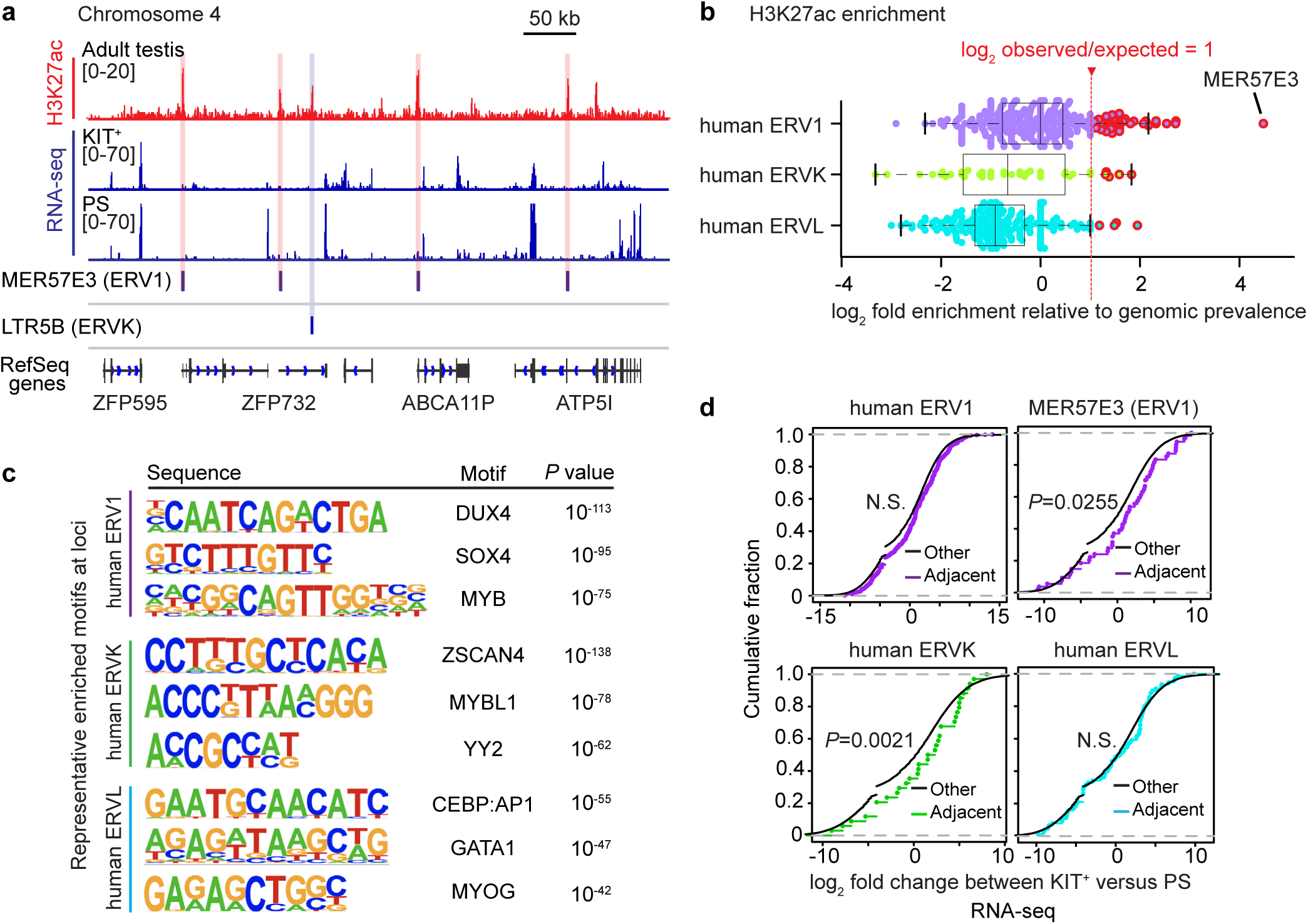
A subset of human ERVKs and ERV1s are associated with meiotic gene expression in humans. (**a**) Representative track view of H3K27ac ChIP-seq in human testis and RNA-seq signals in human KIT^+^ spermatogonia and pachytene spermatocytes (PS). Red and blue rectangles indicate enhancer-like ERV1 and ERVK loci that overlap H3K27ac deposition. (**b**) Beeswarm plots overlaying box and whisker plots; the plots depict the log2 fold enrichments of specific types of ERVs in MACS-identified H3K27ac peak regions in adult human testis, relative to genomic prevalence. Central bars represent medians, the boxes encompass 50% of the data points, and the whiskers indicate 1.5 x interquartile ranges of the data points. Significantly enriched ERV elements were defined as those with values ≥1 log_2_ observed/expected and were outlined in red. (**c**) Motif analyses of enhancer-like human ERV elements for putative transcription factor (TF)-binding sites as called by HOMER (see Methods). (**d**) Cumulative distribution plots of log_2_ fold changes of RNA-seq data between KIT^+^ differentiating spermatogonia and PS for genes adjacent to ERV1s (purple), ERVKs (green), and ERVLs (cyan) as detected by HOMER, all with respect to other expressed genes (black). Significant differences in expression patterns between adjacent genes and other expressed genes were tested via the Wilcoxon rank-sum test. N.S.: not significant.

Next, we sought to test the hypothesis that genes neighboring H3K27ac-enriched ERV loci (>2-fold enrichment) are associated with genes activated after the mitosis-to-meiosis transition (i.e., active in PS compared to KIT^+^ spermatogonia) in humans. In evaluating this possibility, we found that, although genes neighboring H3K27ac-enriched ERV1s did not manifest significant gene expression changes after the mitosis-to-meiosis transition (Fig. 6d), genes adjacent to MER57E3s were significantly activated in PS (Fig. 6d). Notably, genes neighboring H3K27ac-enriched ERVKs tended to be associated with genes activated after the mitosis-to-meiosis transition compared to other genes in the human genome, while genes adjacent to H3K27ac-enriched ERVLs did not show such an association (Fig. 6d). These results suggest that a subset of ERVKs and ERV1s—particularly MER57E3—act as enhancers to activate meiotic genes in humans. Therefore, our results support the notion that ERVK-driven meiotic enhancers are a general feature of mammals, and a subset of ERV1s also functions as enhancers for meiotic genes in humans. Together, we propose that ERV enhancers represent a general mechanism for divergence of transcriptomes during mammalian late spermatogenesis.

## Discussion

In this study, we showed that ERVs act as species-specific enhancers of gene expression after the mitosis-to-meiosis transition in the male germline. Our study identified a novel function of ERVs as enhancers in the germline—distinct from the well-established functions of ERVLs as LTR promoters that drive lncRNA expression^20^. Curiously, over 15% of all oocyte transcripts start at LTR promoters that belong to the ERVL class; these ERVL promoters function during the oocyte-to-embryo transition^46-48^. In contrast to these studies, our study shows that ERVKs function as enhancers in mice and humans, and that a subset of ERV1s act as enhancers in humans. Notably, enhancer-like ERVKs were largely found in intergenic regions (Supplementary Fig. 2) and displayed relatively little enrichment of the promoter mark H3K4me3 (Fig. 3). These novel functions of ERVKs and ERV1s contrast with yet another established function of ERVLs: After fertilization, ERVLs are derepressed and expressed in preimplantation embryos, an essential event in early development^48-50^. Together, our results further expand the repertoire of ERV functions: ERVs are also rapidly evolving enhancers in the germline. Although our analyses focus on the genes adjacent to ERV loci as the targets of enhancer-like ERVKs, there are likely to be many more target genes because long-distance chromatin interactions were found throughout the genome in spermatogenesis^22,23,51^. Thus, further investigation is warranted to identify the full repertoire of genes regulated by enhancer-like ERVs.

Previous studies demonstrated that the transcription factor A-MYB/MYBL1 is a master regulator of the male germline, required for essential events such as meiotic progression and piRNA biogenesis, and for male fertility in general^25,26^. In the present study, we found A-MYB binding sites within enhancer-like ERVKs in mice, and activation of ERV-adjacent genes are regulated by A-MYB. Furthermore, A-MYB binding sites are found in ERVKs and ERV1s in humans. Therefore, retrotransposition of ERVs could have provided new binding sites for key transcription factors to function as newly evolved cis regulatory elements for many genes simultaneously. Importantly, in our accompanying study, we determined that A-MYB is associated with super-enhancers—another type of active enhancers—to drive the expression of key germline genes (Maezawa et al., co-submitted). Therefore, an A-MYB-dependent mechanism appears to lie at the heart of two distinct enhancer types: (1) super-enhancers, which drive the robust activation of germline genes, and (2) ERV-driven, rapidly evolving enhancers, which fine-tune species-specific germline genes. Together, these findings raise an important and intriguing follow-up question: How are A-MYB binding sites on ERVs protected from activation prior to the mitosis-to-meiosis transition? One compelling possibility involves the function of Krüppel-associated box domain zinc finger (KRAB-ZFP) proteins, a family of proteins that have coevolved with ERVs to suppress ERV expression, the consequence of an evolutionary arms race between ERVs and the host genome ^52,53^.

Another important aspect of enhancer-like ERVs is species-specific gene regulation. A recent study reported that, in trophoblast stem cells, RlllLTR13D5s, which comprise a mouse-specific ERVK family, have enhancer functions to establish a trophoblast stem cell-specific regulatory network, and the same study predicted the existence of enhancer-like ERVs in testes and embryonic stem cells^35^. Curiously, placenta is known to be a fast-evolving organ in which many ERVs have been co-opted ^54,55^. Thus, it is intriguing to speculate that ERVs are drivers of species-specific transcriptomes in rapidly evolving organs such as the testis and placenta, although mechanisms underlying intrinsic ERV activity in testes and placenta remain undetermined. Because ERV-based molecular mechanisms expose nuclei to risks of transposition and mutagenesis, their presence and, indeed, apparent importance in germline development is highly enigmatic. If KRAB-ZFP proteins are involved in the control of such mechanisms, it will be crucial to determine the cross-talk between KRAB-ZFP proteins and other means of epigenetic silencing, such as DNA methylation and the piRNA pathway, to understand the precise control of both the silencing of TEs along with their vital activities in the germline.

## Accession Codes

H3K27ac ChIP-seq data reported in this study are described in the accompanying study (Maezawa et al., co-submitted) deposited to the Gene Expression Omnibus (GEO) under the accession number GSE130652. The following secure token has been created to allow review of record GSE130652 while it remains in private status: sbslqciwldgxpof.

## Acknowledgements

We thank Matthew Weirauch and members of the Namekawa laboratory for discussion and helpful comments regarding the manuscript, and the laboratory of Bradley Bernstein for providing human testis H3K27ac ChIP-seq data (ENCSR136ZQZ, ENCODE). Funding sources: Lalor Foundation Postdoctoral Fellowship to A.S.; The research project grant by the Azabu University Research Services Division, Ministry of Education, Culture, Sports, Science and Technology (MEXT)-Supported Program for the Private University Research Branding Project, (2016-2019), Grant-in-Aid for Research Activity Start-up (19K21196), and The Uehara Memorial Foundation Research incentive grant (2018) to S.M.; Albert J. Ryan Fellowship to K.G.A.; National Institute of Health (NIH) DP2 GM119134 to A.B.; March of Dimes Prematurity Research Centre Collaborative Grant (#22-FY14-470) to M.P.; and NIH R01 GM122776 to S.H.N.

## Author contributions

The manuscript was written by A.S., K.G.A., and S.H.N., with critical feedback from all other authors, and A.S. and S.H.N. designed the study. S.M. performed the H3K27ac ChIP-seq experiments. A.S. performed immunostaining. A.S., K.G.A., M.Y., A.B., M.P., and S.H.N. designed and interpreted the computational analyses; A.S. performed the majority of computational analyses. S.H.N. supervised the project.

## Methods

### Animals

Mice were maintained and used according to the guidelines of the Institutional Animal Care and Use Committee (protocol no. IACUC2018-0040) at Cincinnati Children’s Hospital Medical Center.

### Data availability

NGS datasets used in this study are publicly available and referenced within the article. All 50 datasets from other resources are listed and referenced in Supplementary Table 3^5,7,21,26,27,32,33,56-60^.

### Expression analyses for repetitive elements

To analyze the expression of repetitive elements in spermatogenesis, a current build of rodent repetitive sequences was downloaded from Repbase^28^ (http://www.girinst.org/repbase/) and then filtered for *Mus musculus* and ancestral (shared) sequences. Using the bowtie2-build function, a bowtie2 index was generated from a FASTA file of 1,195 mouse repetitive elements. Raw RNA-seq reads were aligned to the indexed repetitive sequence dataset using bowtie2 version 2.3.3.1 with default settings^61^. Aligned reads for each repetitive element were quantified with samtools v 1.5 idxstats^62^ to calculate RPKM values. Differential expression analysis was performed with edgeR^29^ (version 3.18.1). Using the trinityrnaseq package^63^ version 2.0.6, the k-means clustering algorithm was used to obtain plots for scaled, normalized relative expression data on a log scale for each expressed repetitive element. Further analysis was performed with base R and visualized as heat maps using Morpheus (Broad Institute).

### RNA-seq analysis

Raw RNA-seq reads were aligned to either the mouse (GRCm38/mm10) or human (GRCh38/hg38) genomes using Hisat2^64^ version 2.1.0, and uniquely aligned reads were extracted by calling grep with the -v option. To quantify uniquely aligned reads on respective annotated transcript loci (NCBI RefSeq transcripts), the htseq-count function, part of the HTSeq package^65^, was used. The RPKM expression levels for each transcript were calculated using StringTie^66^ version 1.3.4. To detect differentially expressed genes between two biological samples, a read count output file was input to the DESeq2 package (version 1.16.1)^67^; then, the program functions DESeqDataSetFromMatrix and DESeq were used to compare each gene’s expression level between two biological samples. Differentially expressed genes were identified through two criteria: (1) ≥2-fold change and (2) binomial tests (*P* adj < 0.01; *P*values were adjusted for multiple testing using the Benjamini-Hochberg method). To perform gene ontology analysis, the functional annotation clustering tool in DAVID (version 6.8)^68^ was used, and a background of all mouse genes was applied. Biological process term groups with a significance of *P* < 0.05 (modified Fisher’s exact test) were considered significant. Further analysis was performed with R (version 3.4.0) and visualized as heat maps using Morpheus (https://software.broadinstitute.org/morpheus, Broad Institute). To visualize read enrichments over representative genomic loci, TDF files were created from sorted BAM files using the IGVTools count function (Broad Institute). Figures of continuous tag counts over selected genomic intervals were created in the IGV browser (Broad Institute).

### ATAC- and ChIP-seq analyses

Raw ATAC- and ChIP-seq reads were aligned to either the mouse (GRCm38/mm10) or human (GRCh38/hg38) genomes using bowtie2 version 2.3.3.1 with default settings^61^; the reads were filtered to remove alignments mapped to multiple locations by calling grep with the -v option. To identify accessible ERV elements during spermatogenesis, we assembled mouse ERV FASTA references for ATAC-seq alignments (see below). Data from replicates were pooled before identifying regions of enriched occupancy via MACS^69^ version 1.4.2 with default arguments; a cutoff *P* value of 10^−25^ was used. The program ngs.plot^70^ was used to draw tag density plots for H3K27ac, H3K4me3, and H3K27me3 enrichment within ± 1 kb of identified enhancer-like ERVKs (see below). Further analysis was performed with R (version 3.4.0) and visualized as heat maps using Morpheus (https://software.broadinstitute.org/morpheus, Broad Institute). To visualize read enrichment over representative genomic loci, TDF files were created from sorted BAM files using the IGVTools count function (Broad Institute). Figures for continuous tag counts over selected genomic intervals were created in the IGV browser (Broad Institute).

### Identification of enhancer-like ERV elements across mouse and human genomes

Annotation per UCSC RepeatMasker Viz tracks (GRCm38/mm10) was used in the initial screening to identify enhancer-like ERVs in mouse spermatogenesis. Interspersed genomic ERV1, ERVK, and ERVL full-length sequences were extracted using the Bedtools^71^ (version 2.26.0) function getfasta. Using the bowtie2-build function, a bowtie2 index was generated from FASTA files. Raw ATAC-seq reads were aligned to indexed ERV sequence datasets using bowtie2^61^ version 2.3.3.1 with default settings. Aligned reads for each ERV element were quantified with the samtools (version 1.5) function idxstats in order to calculate RPKM values. To determine significant differences in read counts, edgeR^29^ (version 3.18.1) was used. To determine statistical significance for differential accessible loci, only specific ERV loci that showed >2 fold changes in expression and *P*-values of <0.05 were considered. The k-means clustering algorithm was used to obtain plots for scaled, normalized relative expression data on a log scale (log2 RPKM+1). The top 3 enriched ERV subfamilies were defined as accessible ERV candidates, ERV1: RLTR23, RLTR41, and RMER5; ERVK: RLTR10, RMER17, and RLTR51; ERVL: ORR1, MTE, and MTE2: (Fig. 2a, b). To confirm these results, a second screening was performed. The top 3 overrepresented ERV families were determined through comparison of the observed numbers of copies from a family overlapping an MACS-defined ATAC-seq peak region against expected background. The background genomic prevalence was estimated by generating the same number of randomized datasets as each ATAC-seq peak. We computed the number of overlapping ERV copies within ATAC-seq peak regions (observation) and background genomic prevalence (expectation) using custom shell scripts that call the Bedtools^71^ (version 2.26.0) function intersect. Specific ERV subfamilies and subgroups evincing ≥2-fold observed/expected enrichment were defined as “enhancer-like” ERVs (Fig. 2c), and these loci were confirmed via analyses of H3K27ac enrichment with the program ngs.plot^70^. To identify human enhancer-like ERVs in spermatogenesis, this method was performed with UCSC RepeatMasker Viz tracks (GRCh38/hg38) and human testis H3K27ac ChIP-seq data (Fig. 6a).

### Detection of adjacent genes and transcription binding motifs for enhancer-like ERVs

To identify the adjacent genes and overlapping genomic features of enhancer-like mouse and human ERVs, we used the HOMER^36^ (version 4.9) function annotatePeaks.pl. The genes identified as those nearest to enhancer-like ERVs were termed “adjacent genes”. Enrichment of known motifs within enhancer-like ERV loci in mouse and human were analysed with the HOMER^36^ (version 4.9) function findMotifsGenome.pl using default parameters and a fragment size of -gain. All known motifs used in this study were defined with HOMER.

### Evaluation of sequence similarities across mammalian species

To calculate sequence similarities and detect orthologous genes adjacent to mouse enhancer-like ERVKs across other mammalian species (that is, Rat, Rabbit, Marmoset, Gorilla, Chimpanzee, and Human), a list of 543 adjacent genes was applied to BioMart^72^ in order to compute sequence similarities (percent identities of target genes in other species comparted to mouse query genes).

### Determining the protein coding probability of ERVs

To evaluate the protein coding potential of ERVs with expression levels of ≤5 RPKM in pachytene spermatocytes, we used the bioinformatics tool Coding Potential Assessment Tool^31^ (CPAT; http://lilab.research.bcm.edu/cpat/). Protein-coding genes (NCBI RefSeq annotated) were used as a control; protein coding potentials were calculated with default settings.

### Histology and immunofluorescence analyses

For preparation of testicular paraffin blocks, testes were fixed with 4% paraformaldehyde (PFA) overnight at 4°C. Testes were dehydrated and embedded in paraffin. For histological analysis, 5 µm-thick paraffin sections were deparaffinized and autoclaved in target retrieval solution (DAKO) for 10 min at 121°C. Sections were blocked with Blocking One Histo (Nacalai) for 1 h at room temperature and then incubated with primary antibodies (α-γH2AX (05-636-AF647, Millipore) and α-MYBL1 (NBP1-90171, Novus Biologicals) overnight at 4°C. The resulting signals were detected by incubation with secondary antibodies conjugated to fluorophores (Thermo Fisher Scientific, Biotium, or Jackson ImmunoResearch). Sections were counterstained with DAPI. Images were obtained via TiE fluorescence microscopy (Nikon) and processed with NIS-Elements (Nikon) and ImageJ (National Institutes of Health).

## References

1. Ramskold, D., Wang, E.T., Burge, C.B. & Sandberg, R. An abundance of ubiquitously expressed genes revealed by tissue transcriptome sequence data. PLoS Comput Biol 5, e1000598 (2009).

2. Brawand, D. et al. The evolution of gene expression levels in mammalian organs. Nature 478, 343–8 (2011).

3. Soumillon, M. et al. Cellular source and mechanisms of high transcriptome complexity in the mammalian testis. Cell Rep 3, 2179–90 (2013).

4. Lambert, S.A. et al. The Human Transcription Factors. Cell 172, 650–665 (2018).

5. Hasegawa, K. et al. SCML2 Establishes the Male Germline Epigenome through Regulation of Histone H2A Ubiquitination. Dev Cell 32, 574–88 (2015).

6. Sin, H.S., Kartashov, A.V., Hasegawa, K., Barski, A. & Namekawa, S.H. Poised chromatin and bivalent domains facilitate the mitosis-to-meiosis transition in the male germline. BMC Biol 13, 53 (2015).

7. Lesch, B.J., Silber, S.J., McCarrey, J.R. & Page, D.C. Parallel evolution of male germline epigenetic poising and somatic development in animals. 48, 888–94 (2016).

8. Waterston, R.H. et al. Initial sequencing and comparative analysis of the mouse genome. Nature 420, 520–62 (2002).

9. Lander, E.S. et al. Initial sequencing and analysis of the human genome. Nature 409, 860–921 (2001).

10. McClintock, B. The origin and behavior of mutable loci in maize. Proc Natl Acad Sci U S A 36, 344–55 (1950).

11. Rebollo, R., Romanish, M.T. & Mager, D.L. Transposable elements: an abundant and natural source of regulatory sequences for host genes. Annu Rev Genet 46, 21–42 (2012).

12. Friedli, M. & Trono, D. The developmental control of transposable elements and the evolution of higher species. Annu Rev Cell Dev Biol 31, 429–51 (2015).

13. Chuong, E.B., Elde, N.C. & Feschotte, C. Regulatory activities of transposable elements: from conflicts to benefits. Nat Rev Genet 18, 71–86 (2017).

14. Garcia-Perez, J.L., Widmann, T.J. & Adams, I.R. The impact of transposable elements on mammalian development. Development 143, 4101–4114 (2016).

15. Thompson, P.J., Macfarlan, T.S. & Lorincz, M.C. Long Terminal Repeats: From Parasitic Elements to Building Blocks of the Transcriptional Regulatory Repertoire. Mol Cell 62, 766–76 (2016).

16. Zamudio, N. & Bourc’his, D. Transposable elements in the mammalian germline: a comfortable niche or a deadly trap? Heredity (Edinb) 105, 92–104 (2010).

17. Crichton, J.H., Dunican, D.S., Maclennan, M., Meehan, R.R. & Adams, I.R. Defending the genome from the enemy within: mechanisms of retrotransposon suppression in the mouse germline. Cell Mol Life Sci 71, 1581–605 (2014).

18. Ku, H.Y. & Lin, H. PIWI proteins and their interactors in piRNA biogenesis, germline development and gene expression. Natl Sci Rev 1, 205–218 (2014).

19. Watanabe, T., Cheng, E.C., Zhong, M. & Lin, H. Retrotransposons and pseudogenes regulate mRNAs and lncRNAs via the piRNA pathway in the germline. Genome Res 25, 368–80 (2015).

20. Davis, M.P. et al. Transposon-driven transcription is a conserved feature of vertebrate spermatogenesis and transcript evolution. EMBO Rep 18, 1231–1247 (2017).

21. Maezawa, S., Yukawa, M., Alavattam, K.G., Barski, A. & Namekawa, S.H. Dynamic reorganization of open chromatin underlies diverse transcriptomes during spermatogenesis. Nucleic Acids Res 46, 593–608 (2018).

22. Alavattam, K.G. et al. Attenuated chromatin compartmentalization in meiosis and its maturation in sperm development. Nat Struct Mol Biol 26, 175–184 (2019).

23. Patel, L. et al. Dynamic reorganization of the genome shapes the recombination landscape in meiotic prophase. Nat Struct Mol Biol 26, 164–174 (2019).

24. Wang, Y. et al. Reprogramming of Meiotic Chromatin Architecture during Spermatogenesis. Mol Cell 73, 547-561.e6 (2019).

25. Bolcun-Filas, E. et al. A-MYB (MYBL1) transcription factor is a master regulator of male meiosis. Development 138, 3319–30 (2011).

26. Li, X.Z. et al. An ancient transcription factor initiates the burst of piRNA production during early meiosis in mouse testes. Mol Cell 50, 67–81 (2013).

27. Kobayashi, H. et al. Contribution of intragenic DNA methylation in mouse gametic DNA methylomes to establish oocyte-specific heritable marks. PLoS Genet 8, e1002440 (2012).

28. Bao, W., Kojima, K.K. & Kohany, O. Repbase Update, a database of repetitive elements in eukaryotic genomes. Mob DNA 6, 11 (2015).

29. Robinson, M.D., McCarthy, D.J. & Smyth, G.K. edgeR: a Bioconductor package for differential expression analysis of digital gene expression data. Bioinformatics 26, 139–40 (2010).

30. Meyer, T.J., Rosenkrantz, J.L., Carbone, L. & Chavez, S.L. Endogenous Retroviruses: With Us and against Us. Front Chem 5, 23 (2017).

31. Wang, L. et al. CPAT: Coding-Potential Assessment Tool using an alignment-free logistic regression model. Nucleic Acids Res 41, e74 (2013).

32. Jung, Y.H. et al. Chromatin States in Mouse Sperm Correlate with Embryonic and Adult Regulatory Landscapes. Cell Rep 18, 1366–1382 (2017).

33. Maezawa, S. et al. Polycomb protein SCML2 facilitates H3K27me3 to establish bivalent domains in the male germline. Proc Natl Acad Sci U S A 115, 4957–4962 (2018).

34. Adams, S.R. et al. RNF8 and SCML2 cooperate to regulate ubiquitination and H3K27 acetylation for escape gene activation on the sex chromosomes. PLoS Genet 14, e1007233 (2018).

35. Chuong, E.B., Rumi, M.A., Soares, M.J. & Baker, J.C. Endogenous retroviruses function as species-specific enhancer elements in the placenta. Nat Genet 45, 325–9 (2013).

36. Heinz, S. et al. Simple combinations of lineage-determining transcription factors prime cis-regulatory elements required for macrophage and B cell identities. Mol Cell 38, 576–89 (2010).

37. Aloisio, G.M. et al. PAX7 expression defines germline stem cells in the adult testis. J Clin Invest 124, 3929–44 (2014).

38. Gely-Pernot, A. et al. Retinoic Acid Receptors Control Spermatogonia Cell-Fate and Induce Expression of the SALL4A Transcription Factor. PLoS Genet 11, e1005501 (2015).

39. Le Bouffant, R. et al. Msx1 and Msx2 promote meiosis initiation. Development 138, 5393–402 (2011).

40. Wu, S., Hu, Y.C., Liu, H. & Shi, Y. Loss of YY1 impacts the heterochromatic state and meiotic double-strand breaks during mouse spermatogenesis. Mol Cell Biol 29, 6245–56 (2009).

41. Young, J.M. et al. DUX4 binding to retroelements creates promoters that are active in FSHD muscle and testis. PLoS Genet 9, e1003947 (2013).

42. Isbel, L. et al. Trim33 Binds and Silences a Class of Young Endogenous Retroviruses in the Mouse Testis; a Novel Component of the Arms Race between Retrotransposons and the Host Genome. PLoS Genet 11, e1005693 (2015).

43. Laiho, A., Kotaja, N., Gyenesei, A. & Sironen, A. Transcriptome profiling of the murine testis during the first wave of spermatogenesis. PLoS One 8, e61558 (2013).

44. Smith, T.B., Baker, M.A., Connaughton, H.S., Habenicht, U. & Aitken, R.J. Functional deletion of Txndc2 and Txndc3 increases the susceptibility of spermatozoa to age-related oxidative stress. Free Radic Biol Med 65, 872–881 (2013).

45. Mueller, J.L. et al. The mouse X chromosome is enriched for multicopy testis genes showing postmeiotic expression. Nat Genet 40, 794–9 (2008).

46. Peaston, A.E. et al. Retrotransposons regulate host genes in mouse oocytes and preimplantation embryos. Dev Cell 7, 597–606 (2004).

47. Veselovska, L. et al. Deep sequencing and de novo assembly of the mouse oocyte transcriptome define the contribution of transcription to the DNA methylation landscape. Genome Biol 16, 209 (2015).

48. Franke, V. et al. Long terminal repeats power evolution of genes and gene expression programs in mammalian oocytes and zygotes. Genome Res 27, 1384–1394 (2017).

49. De Iaco, A. et al. DUX-family transcription factors regulate zygotic genome activation in placental mammals. Nat Genet 49, 941–945 (2017).

50. Hendrickson, P.G. et al. Conserved roles of mouse DUX and human DUX4 in activating cleavage-stage genes and MERVL/HERVL retrotransposons. Nat Genet 49, 925–934 (2017).

51. Wang, M. et al. Single-Cell RNA Sequencing Analysis Reveals Sequential Cell Fate Transition during Human Spermatogenesis. Cell Stem Cell 23, 599-614.e4 (2018).

52. Ecco, G. et al. Transposable Elements and Their KRAB-ZFP Controllers Regulate Gene Expression in Adult Tissues. Dev Cell 36, 611–23 (2016).

53. Imbeault, M., Helleboid, P.Y. & Trono, D. KRAB zinc-finger proteins contribute to the evolution of gene regulatory networks. Nature 543, 550–554 (2017).

54. Dunn-Fletcher, C.E. et al. Anthropoid primate-specific retroviral element THE1B controls expression of CRH in placenta and alters gestation length. PLoS Biol 16, e2006337 (2018).

55. Chuong, E.B. The placenta goes viral: Retroviruses control gene expression in pregnancy. PLoS Biol 16, e3000028 (2018).

56. Li, D. et al. Chromatin Accessibility Dynamics during iPSC Reprogramming. Cell Stem Cell 21, 819-833.e6 (2017).

57. Cao, S. et al. Chromatin Accessibility Dynamics during Chemical Induction of Pluripotency. Cell Stem Cell 22, 529-542.e5 (2018).

58. He, S. et al. Hemi-methylated CpG sites connect Dnmt1-knockdown-induced and Tet1-induced DNA demethylation during somatic cell reprogramming. Cell Discov 5, 11 (2019).

59. Guo, J. et al. Chromatin and Single-Cell RNA-Seq Profiling Reveal Dynamic Signaling and Metabolic Transitions during Human Spermatogonial Stem Cell Development. Cell Stem Cell 21, 533-546.e6 (2017).

60. An integrated encyclopedia of DNA elements in the human genome. Nature 489, 57–74 (2012).

61. Langmead, B. & Salzberg, S.L. Fast gapped-read alignment with Bowtie 2. Nat Methods 9, 357–9 (2012).

62. Li, H. et al. The Sequence Alignment/Map format and SAMtools. Bioinformatics 25, 2078–9 (2009).

63. Grabherr, M.G. et al. Full-length transcriptome assembly from RNA-Seq data without a reference genome. Nat Biotechnol 29, 644–52 (2011).

64. Kim, D., Langmead, B. & Salzberg, S.L. HISAT: a fast spliced aligner with low memory requirements. Nat Methods 12, 357–60 (2015).

65. Anders, S., Pyl, P.T. & Huber, W. HTSeq--a Python framework to work with high-throughput sequencing data. Bioinformatics 31, 166–9 (2015).

66. Pertea, M. et al. StringTie enables improved reconstruction of a transcriptome from RNA-seq reads. Nat Biotechnol 33, 290–5 (2015).

67. Love, M.I., Huber, W. & Anders, S. Moderated estimation of fold change and dispersion for RNA-seq data with DESeq2. Genome Biol 15, 550 (2014).

68. Huang da, W., Sherman, B.T. & Lempicki, R.A. Bioinformatics enrichment tools: paths toward the comprehensive functional analysis of large gene lists. Nucleic Acids Res 37, 1–13 (2009).

69. Zhang, Y. et al. Model-based analysis of ChIP-Seq (MACS). Genome Biol 9, R137 (2008).

70. Shen, L., Shao, N., Liu, X. & Nestler, E. ngs.plot: Quick mining and visualization of next-generation sequencing data by integrating genomic databases. BMC Genomics 15, 284 (2014).

71. Quinlan, A.R. & Hall, I.M. BEDTools: a flexible suite of utilities for comparing genomic features. Bioinformatics 26, 841–2 (2010).

72. Durinck, S., Spellman, P.T., Birney, E. & Huber, W. Mapping identifiers for the integration of genomic datasets with the R/Bioconductor package biomaRt. Nat Protoc 4, 1184–91 (2009).

