## Supplemental Figures for "Endogenous retroviruses drive species-specific germline transcriptomes in mammals"

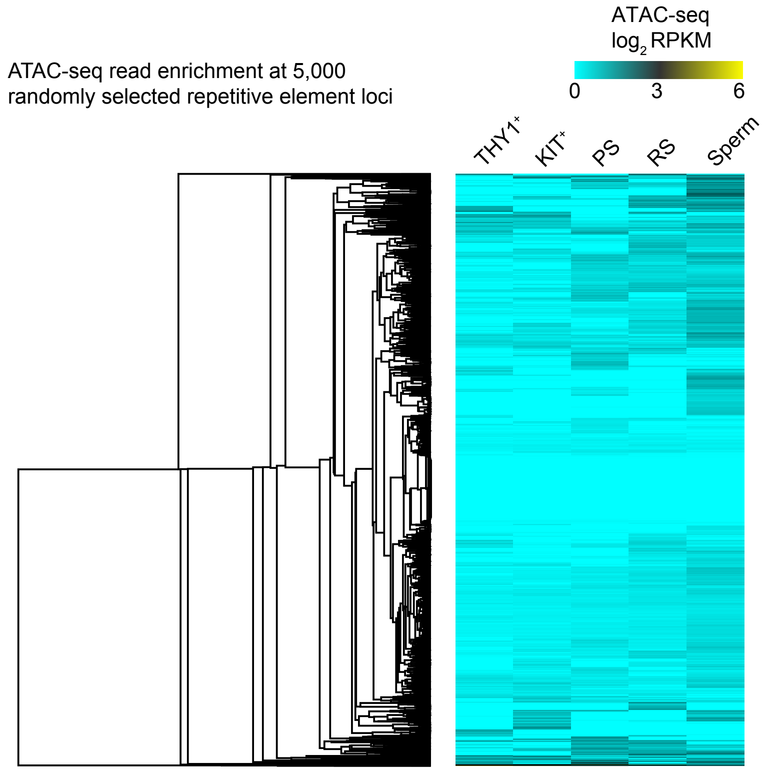


**Supplementary Figure 1. ATAC-seq read enrichment at 5,000 randomly selected repetitive element loci.** Heat map depicting RPKM-normalized ATAC-seq reads at 5,000 randomly selected repetitive element loci during spermatogenesis. $THY1^{+}$, undifferentiated spermatogonia; $KIT^{+}$, differentiating spermatogonia; PS, pachytene spermatocyte; RS, round spermatids; Sperm: epididymal spermatozoa.

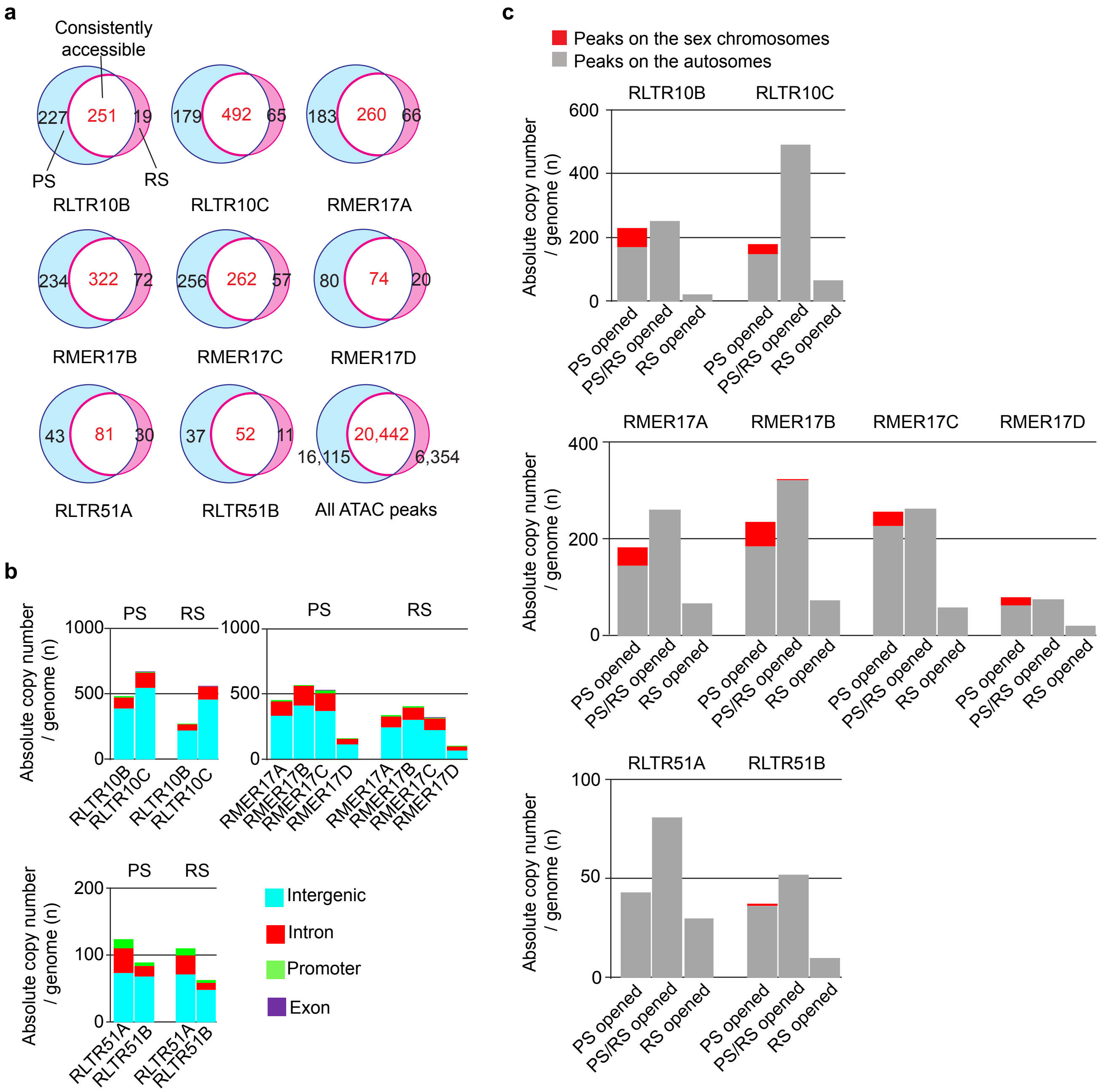


**Supplementary Figure 2. Genomic features of accessible ERVKs in late spermatogenesis.**
(**a**) Venn diagram showing the overlap of young ERVK loci in accessible chromatin between pachytene spermatocytes (PS; blue) and round spermatids (RS; pink). (**b**) Bar charts depicting genomic distributions of young ERVK loci in accessible chromatin. Distributions are shown with colored bars. (**c**) Bar charts indicating absolute copy numbers on autosomes (gray) and sex chromosomes (red) of young ERVKs.

**a** RLTR10B (RepBase consensus)

Similarity: 172/317 (54.26%) by EZBioCloud Pairwise nucleotide sequence alignment for taconomy

```
Mouse RLTR10B: TGTGGGAAGC---CACATGTGCCGTTGCAGGGTGGCGCTGGCTACCGCTGGCCACCACGCATACATAGGCA
Rat RLTR10B:   TGTGGGGAGCGGGTGTAGCGGCAATCCCAAGATGGCGCCCGGGACTGCTGCCGA-----------------

GTAAAGTTTTTTTGCCAAGATGAGGTTTTGAGAATTAACCAAAATTAACCAATCAGATGAGAGACAAGTTAACCAATCA-GAT
-----------GTCTTACGACTAGCACCTGAAAGTAAGCCA----CGCCCCATC--GTGAGAGCTGCGCAGGCGCACCATGAC

GAGAGACAAGTTAACCAATCAGATGTGAGGCATGCAAATGAGGTGGTAAGCATAACCCATGCATAACCAATCCGGGTGTGAGAC
GATAGA----TCAGGCCATGTGACGTGGACCTATGAACGGAGGT--TACGCAGA----CTGGACGGTTGAGGCGGAGTTGAG--

ACGCCTCTCCTAGGCCTATATAAG---CAGCACCAGTTCTGGGGCTTGGGGGTCTCTTCGCCTCTGCAATC---AAGCTCTCC
------------GGAGGTTATAAGGGAGTGCGCTCGGGGGCTGGATTGAGAGATTGTTGCTGCTTGCATGCTAAAAGGTTCCT

CAATAAAC----GTGTGCAGAAGGATCCTGTTGCAGCTGTCGTTCTTGCTGGCGAGTCAGGGCGCGCGCAA
GAATAAACTGCTTTGAGAAGAACG-TCGTGGTGTCGCTCCTTTCCTGCTGGTCGGGGTTGGAAGCGACA--
```

**b** RMER17A (RepBase consensus)

Similarity: 189/319 (59.25%) by EZBioCloud Pairwise nucleotide sequence alignment for taconomy

```
Mouse RMER17A:TGTGGGGAATTGCAGGCTGGTTCCAGTTGAGCTGAGGTCTGAACCCCAGTGGTGATAATTCACCTACATGAC
Rat RMER17A:  ------------------------------------------------------------------------

ACGGTAGGCATTCCCTCATGCTCCTGGAACTCTGGCTCCTGCCTAAGTTACCGCCCCCCACAGCCCCCACAAGAGAAGCAT
-----------------------------------------------------------------------------------

GGTTAGTAGTCACGCAGGCAATGTCCCAAGCTTCTGACCTTCAGGCTAGACTCCTCCC---CAGTTACCTAGCAACGTAAAG--
------------------------------TGTTAGCATTCTGTCTAAGCTCCACCCCCACAGTTACCTGGCAACAGCCAGGT

---------ACCATAAGAGGGGCTGCTCGGCCCCTCCTCGCTCTCTTACTTGTCTCTTCTCTTACTCCTTTGTTCTCTTAACT
ATGCCTGACACTATAAAAGGGGCTGCTTGCCCCCTCCTCACTCTCTTGC---TCTTGCTTCTTGCTCTCTTGCTCTTCCCCTC

TCTCACACTTCTTTCTCCTCTTCCCTTTCTCTTTGTCTTCTCTCCTCTCTCCTTTCTCTCTACTCTTCTCTCAGCCTTCTCTCT
CTTCCCCTTTGTCCCTTCTCTCCCCATTCCC---CTCCCCCCTTCCCTCCACGTGCTCATGGCCGGCCTCT---ACTCCTCTCC

CTCTCTCCCTCTCCCTCCCTCTCTCCTCTCTTATACCCCTGCCTTTCTACAATAAAGCTCTAAAACCATAGAGAGTCTCTGCT
TCTTCTACTCTTCTCTCTCTCGTCCCTCTCCCTTGTCTCGTCCTTT---CATTAAACCTTTCCA-----------CGTGGAAC

CATCAAGATCTGCTGGGCTCACTCTCGTCAGTGTTGGGAACCTCTTCCCCCATCCTTCTCTCCCATAACCCTGGTGGCTTTAGA
CATGTTGGCCTGGTGTGGTTTGTCCGGATGCGAGCCGAGATTTCTACCCCAACA-----------------------------

AAGTAGCTCTGGGGCCCCCAGGTAGGGCTGCCCCTTGGCCACCCCCCGAAGAGTGGGTCAGAGGCTTAGATGCCCACCCAGGG
-----------------------------------------------------------------------------------

ATGAGTGGAAGGTAGATAGCAGCCCTCCCACCTGACTGACCAGAGAACAGGTGGAACTCTGGCGGGGTGTGGGTCTTCCCCTT
-----------------------------------------------------------------------------------

CCCCCTCTTCCCCAGGGCCCCCCACCTTTTTGCTCCCAA
---------------------------------------
```

**Supplementary Figure 3. Sequence similarities of young ERVK elements between mouse and rat.**
(**a**,**b**) Pairwise alignments of RLTR10B (**a**) and RMER17A (**b**) entire consensus sequences between mouse and rat.

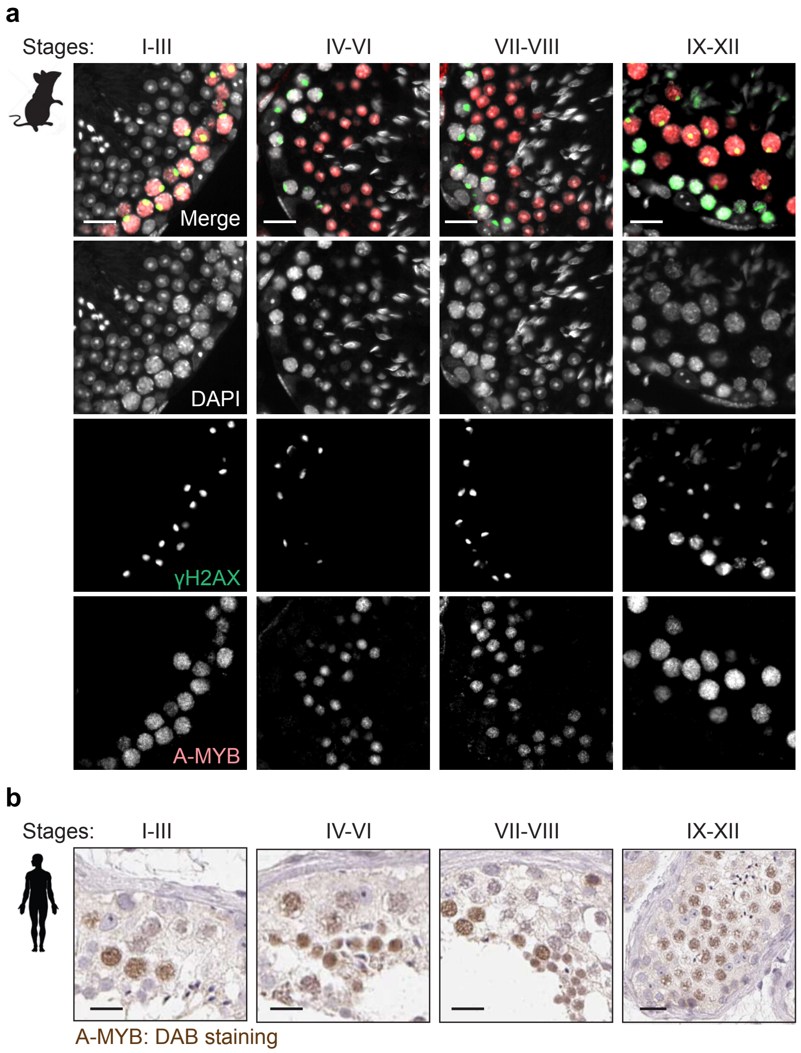


**Supplementary Figure 4. A-MYB are highly expressed both in mouse and human spermatocytes.**
(**a**) Immunofluorescence co-staining of 12-week-old mouse testicular cross sections for A-MYB (red) and γH2AX (green), and counterstaining with DAPI (gray). The Roman numerals indicate cycles of seminiferous epithelium. (**b**) Representative images of immunohistological staining of 29-to-65-year-old human testicular sections for A-MYB (brown) and counterstaining with hematoxylin (adapted from the Human Protein Atlas (www.proteinatlas.org/ENSG00000185697-MYBL1/tissue/testis#img)). Scale Bars: 20 μm.
